# Comparative forewing ultrastructure of high-frequency singing crickets (Gryllidae, Eneopterinae)

**DOI:** 10.1101/2024.12.01.625810

**Authors:** L. Royer, T. Gaiddon, T. Robillard

## Abstract

Male crickets belonging to the tribe Lebinthini (Eneopterinae) produce high-frequency, sometimes ultrasonic, calling songs characterized by a dominant harmonic frequency -a phenomenon known as harmonic hopping. While cricket sound production is well understood, the mechanism allowing the dominant frequency to move to the harmonics of the spectrum is little studied. The wing region known as the harp corresponds to the primary resonator of the cricket tegmina. Here we hypothesize that the surface ultrastructure of this region could play a role in the physical properties of the wings and in their vibratory behavior. In this study, we used scanning electron microscopy to explore the diversity of wing membrane ultrastructure in the harp region of 33 species of Eneopterinae, including species producing both low-frequency and high-frequency calling songs. We highlighted a great diversity of ultrastructure within the subfamily. We defined and measured 5 morphological traits concerning the hexagonal cells visible on the dorsal face of the wing, and the microtrichia, filiform sensory structures present on the ventral face. Significant differences are found in hexagonal cell size and density between males and females, and between the males of the species of the Lebinthini tribe and the non-Lebinthini species. These results suggest a link between the presence of these hexagonal structures and the song frequency value. Surfacic structures could play a role in the stiffening of the wing, a property directly related to vibration frequency of the wing. The observation of these structure and more precisely their size and distribution between high-frequency songs producing species and Low-frequency species constitute a new avenue for the understanding of harmonic-hoppings in Lebinthini crickets.

## Introduction

Acoustic signals are widely used for communication in animals, especially for finding mating partners (e.g., Bradbury & Vehrencamp, 2011; Gerhardt & Huber, 2002). Studying these signals in the context of evolutionary research is important because of their implication in multiple selection pressures involving both sexual selection (species recognition, choice of partner) and natural selection (attraction of predators and parasites, propagation in habitat, competition for acoustic space). In most species using acoustic communication, the male individual is the one emitting the signal via an advertisement call or a calling song, while the female plays the role of receiver. Sound production mechanisms are diverse among the animal kingdom, and many groups possess specialized organs for sound production. In insects, specialized organs and sound production mechanisms are multiple and diverse. The most known cases are cymbalisation in Cicadas and stridulation in Orthopterans (Bennet-Clark, 1999a). While Caeliferans (grasshoppers) usually stridulate by rubbing their femora against their tegmina (sclerified anterior wings), Ensiferans (crickets and katydids) acoustic structures are located on the forewings only (Fig. 1). Regardless of the sound production mechanism, the signal is composed of multiple dimensions such as rhythm, intensity and frequency. The call frequency spectrum is usually composed of a dominant peak that usually corresponds to the carrier (fundamental) frequency, followed by a series of harmonics. Harmonic frequencies are integer multiples of the carrier frequency and of decreasing intensity (Speaks, 1996). In some taxa, such as *Rhinolophus philippinensis* bats (Kingston & Rossiter, 2004) or *Selasphorus* hummingbirds (Clark, 2014), a singular phenomenon is observed, leading the dominant frequency of the calling song to be one of the harmonics, and not the carrier anymore. This phenomenon is referred to as harmonic hopping (Clark, 2014; Kingston & Rossiter, 2004; Tan et al., 2021) and results in multiplying the dominant frequency value (doubled if the second harmonic becomes dominant, tripled if this is the third, etc.). Such changes of the call frequency can have a strong impact on the signal properties, by potentially affecting interactions with predators, parasites and competitors (Robillard & Desutter-Grandcolas, 2004).

**Fig 1:**
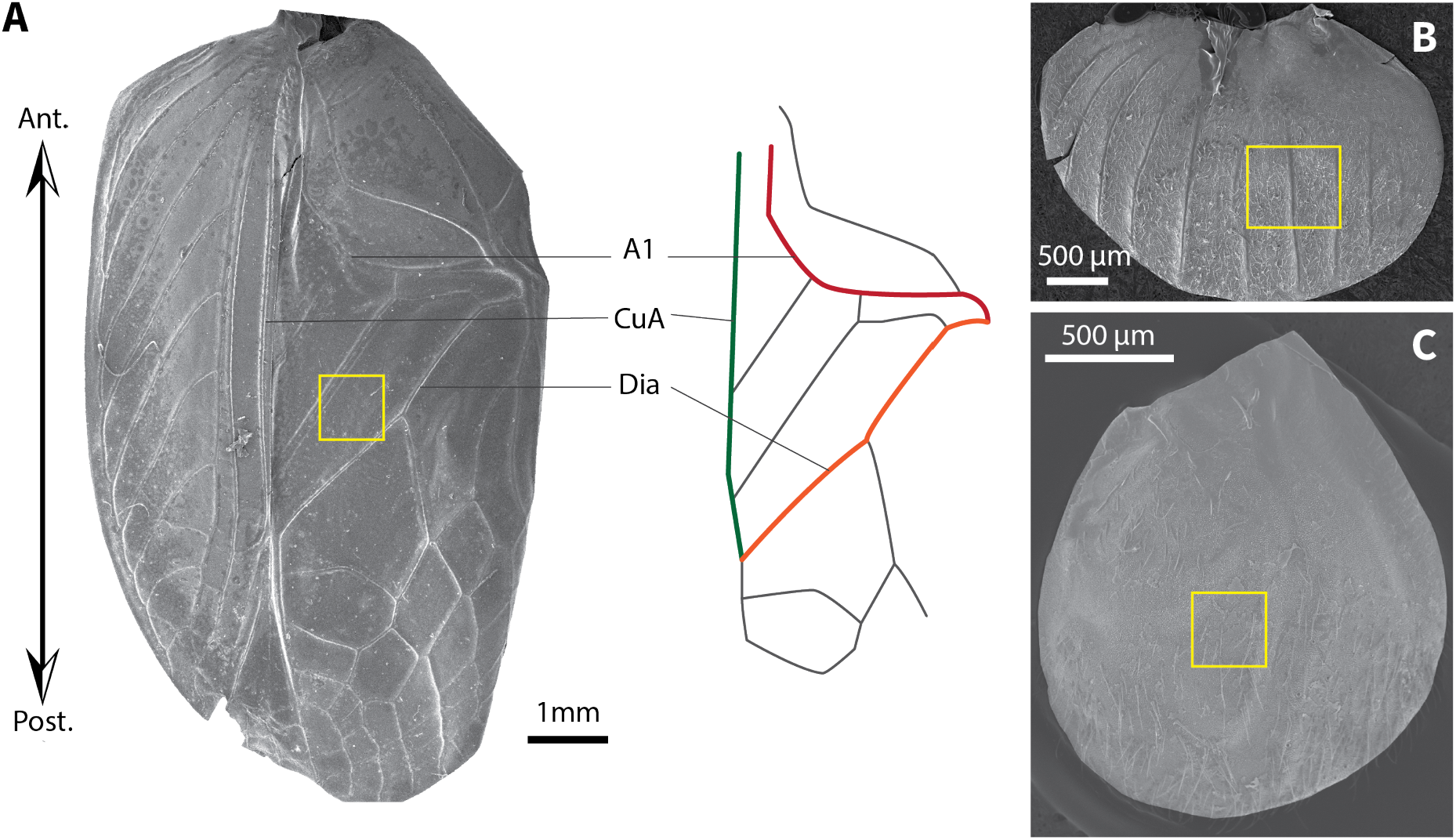
**A**) SEM picture of the left forewing of *Nisitrus danum* male, with simplified venation pattern on the left. The three colored veins delimit the harp. 1A = first anal vein carrying the stridulatory file, CuA = anterior cubital vein, Dia = diagonal vein. **B**) SEM picture of the left forewing of *Agnothecous obscurus* female. **C**) SEM picture of the left wing bud (last larval stage) of *Gnominthus baitabagus*. Yellow rectangles show the sampled area.

Although dominant harmonics are common in multiple animal groups (frogs, bats, birds, insects), harmonic hopping has been studied only in a few clades. In particular, harmonic hoppings are well known in crickets of the subfamily Eneopterinae Saussure, 1874 (Robillard et al., 2007, 2013; Robillard & Desutter-Grandcolas, 2004; Tan et al., 2021). Numerous events of harmonic hoppings happened during the evolutionary history of this clade (Tan et al., 2021) and are supposed to be associated with the diversification of the tribe Lebinthini Robillard, 2004 (Robillard & Desutter-Grandcolas, 2004). Most of the 175 species of this tribe use high-frequency signals acquired from one or more harmonic hopping events (Tan et al., 2021), with subsequent diversification both in terms of number of species and of frequency spectra. These high-frequency signals are also associated with a distinctive communication system: Lebinthini females lack phonotaxis and they respond to male calls by producing a specific vibrational signal (Hofstede et al., 2015).

The sound production mechanism of crickets includes two distinct phases: 1) the initial vibration is generated during the stridulation phase from the fast scraping of the stridulatory file, then 2) the sound is amplified by resonance of the forewings membrane during the amplification phase (Bennet-Clark, 1999a; Michelsen, 1998). For most cricket species, the principal resonator corresponds to the harp region, a triangle-shaped area posterior to the file, in the dorsal field (Michelsen, 1998; Montealegre-Z et al., 2011). Cricket songs usually show a relatively low fundamental frequency, ranging between 2 and 8 kHz.

Previous studies on the stridulatory mechanisms of crickets revealed that some morphological characteristics such as the length of the file, inter-teeth distances and speed of wing closing movement, have a direct influence on the carrier frequency of the song (Montealegre-Z, 2009; Robillard et al., 2013). However several studies more specifically on Lebinthini species suggested that the amplification of harmonics frequencies correspond to modifications of the amplification phase, through changes of the resonator properties (Robillard et al., 2013, 2015).

Physical properties of the wing membrane, such as elasticity and stiffness can strongly contribute to determine the frequency of the sound generated during stridulation (Sample et al., 2015, Gaiddon et al. submit). These properties depend both on the wing shape and dimensions (Gromougin et al. in prep.), and on the material making up the wings (Michels et al., 2016, Gaiddon et al. submit). Among the factors possibly involved, the surface ultrastructure of the wing membrane is also likely to play a role in modifying the wings’ physical properties. Few studies have investigated the cuticular structures of crickets. In a study describing the structures of the ventral and dorsal surfaces of the tegmina of various species from different clades of the superfamily Grylloidea, Desutter-Grandcolas (1995, 1998) suggested that hexagonal shaped ultrastructures of the wings membrane could be related to vibrational characteristics. Other studies have focused on the microscopic structure of the stridulatory file (Li et al., 2016), or on the microscopic structures of the body (Barranco & Molina-Pardo, 2021). No study has yet been conducted to compare wing ultrastructure among closely related species within a phylogenetic framework. Similarly, the supposed correlation between song parameters and tegminal ultrastructure has never been formally tested using a comprehensive sampling of species.

Here we aim to explore the diversity of wing ultrastructure at the harp region in males and females for 33 species of the Eneopterinae subfamily using scanning electron microscopy. We expect to observe micro-structures similar to those observed in previous studies (Desutter-Grandcolas, 1995; Eisenbeis & Wichard, 2012) with disparity in conformations across the subfamily. If a link between ultrastructural traits and sound production exists, we expect to find intraspecific sexual differences, as well as differences between species producing high-frequency calls and the others. To test this hypothesis, we measured the density and size of the two main ultrastructures highlighted in other crickets clades: hexagonal cells on the dorsal face of the forewings, and microtrichia on the ventral face. We performed mean comparisons between pairs of males and females and between acoustic groups.

To also investigate the development and origins of the observed structures, we imaged the wing buds of the juveniles for a subsample (6) of species. We expect to observe the same developing structures as in adult wings, with differences between the juvenile stages, in order to test the hypothesis that the hexagonal structures and microtrichia develop during the last stage of development in the last wing bud.

## Materials and methods

### Taxonomic sampling

We sampled 33 species of the cricket subfamily Eneopterinae for ultrastructural observations. The sampled species represent 16 genera within the 5 extant tribes of these crickets and thus represent the acoustic and phylogenetic diversity of the subfamily (Tan et al., 2021; Vicente et al., 2017). While cricket wings are almost symmetrical, their functioning is asymmetrical, and previous studies have shown that the left wing tends to vibrate more than the right one in some species (Montealegre-Z et al., 2011)Gaiddon et al submit, in particular in eneopterines (Robillard et al., 2013; Jonsson et al. submit). Therefore, after checking in a few samples that ultrastructural elements were equivalent between the two wings, we focused our observations on the left forewing of male and female specimens. For each species and sex, two left forewings have been collected from two adult specimens from the collections of the Muséum national d’Histoire naturelle (MNHN), in order to study both ventral and dorsal faces of the wing. For six species (4 Lebinthini and 2 Nisitrini), we also studied wing buds from the two last juvenile stages, for both sexes and in ventral and dorsal views. Most of these juvenile samples have been collected on living specimens from cricket colonies maintained in MNHN, completed by juvenile samples from the collections when necessary.

### Scanning electron microscopy

Pinned specimens were rehydrated for 30 minutes using small pieces of humidified cotton, and their left wing were cut and glued on cardboard holders. Samples were then mounted on stubs and platinum-coated (8 nm) with a Leica EM ACE600 metallizer. Observations were made under two scanning different electron microscopes (SEM): Hitachi SU3500 SEM or a FEG TESCAN SEM, both from the UAR 2AD microscopy service of the MNHN at magnifications up to x2000.

For male samples, SEM pictures were taken in the same area of interest for each face of the wing corresponding to the middle of the harp **(Fig. 1A)**. Female venation being very different, pictures were taken in areas corresponding to the location of the harp in males **(Fig. 1B)**. Similarly, for the juvenile wing buds, as venation is not yet in place, pictures of the membrane were taken to best represent the whole wing bud surface **(Fig. 1C)**. Regardless of the sample, a minimum of 3 images were taken under different magnifications, starting with broad views of the harp region and its surroundings, followed by detailed observation at higher magnifications.

### Measurements and statistical analyses

Based on the morphological exploration of the surfacic features of the wings, we defined 5 traits concerning 2 types of structures previously observed in crickets: filamentous structures observed on the ventral face, referred to as microtrichia, and hexagonal cells observed on the dorsal face of the forewing (Desutter-Grandcolas, 1995). All the measurements were made from the SEM images using the program FIJI (Schindelin et al., 2012) for the traits defined as follows:

- The density of microtrichia on the ventral face of males, females and juvenile wings was semi-automatically counted within a square of 0.07±0.03 mm^2^ using the Particles analysis function of the software.
- The microtrichia length in males and females, was measured as the mean of 30 measurements from the same picture. The length is considered as null in the absence of microtrichia.
- The density of hexagonal-shaped structures on the dorsal face was manually measured in males and females by counting the number of structural units at x500 magnification within a square of 0.05±0.01 mm^2^.
- The hexagonal structure size was measured as the mean diameter of 30 structures from the same picture in males and females. The diameter is 0 in the absence of the structures
- The thickness of the hexagonal structure walls was measured in both males and females as the mean of 30 manual measurements per sample. In species presenting very thin flat structures (scale-like rather than hexagonal cells), the walls thickness was considered as null.

Wilcoxon Mean comparison tests with paired samples (males and females of the same species being paired) were performed using the wilcox.test function from Rbase with R version 4.2.2 (R Core Team, 2022).

To test the effect of size on the ultrastructural traits, we used the mean pronotum length (in mm) per species as representing the body size measure and mean harp area (in mm^2^) as harp size (measured from a previous study: Grosmougin et al., in prep). The same dataset has been used for females and males, as size dimorphism is negligible in Eneopterinae. Pearson’s correlation tests were performed using the cor.test function from Rbase.

## Results

### Adult forewings ultrastructure

SEM observations of the forewing surfaces show a great diversity of ultrastructural conformations among the Eneopterinae crickets. At very high magnification, most of the samples exhibited a microscopic granularity on both the ventral and dorsal surfaces of the wing (**Fig. 2**,**3**,**4**,**5** right panels) in both sexes. In 3 samples: *Falcerminthus sandakan* male dorsal face (**Fig. 2C**), *Nisitrus malaya* female dorsal face (**Fig. 3E-F**) and *Xenogryllus eneopteroides* female ventral face (**Fig. 5B-C**), the granularity was difficult to distinguish, but it remains visible on the other side of the forewing, which allows us to hypothesize that this granularity is present in all the studied species.

**Fig 2:**
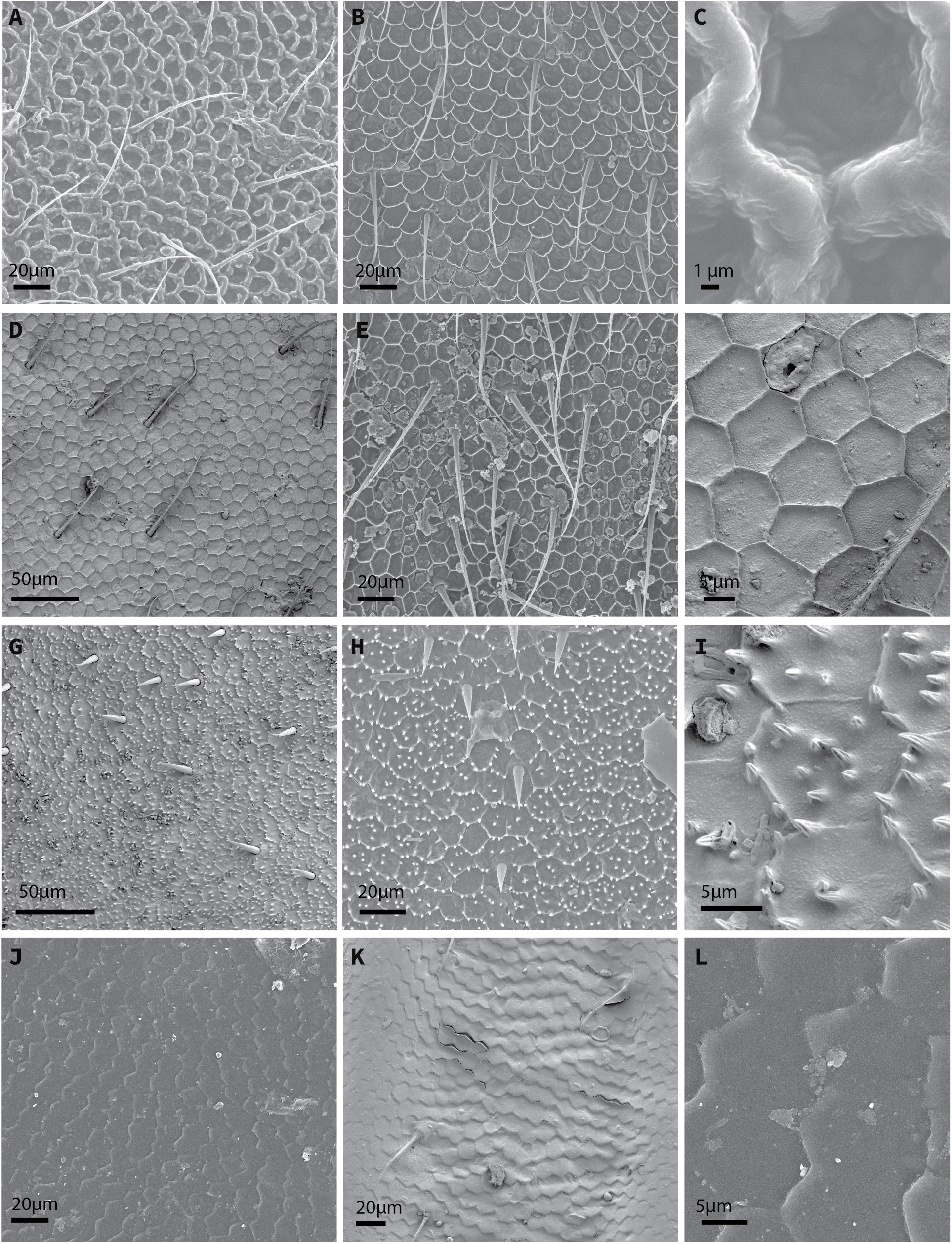
SEM pictures of the dorsal face of the forewing membranes ultrastructure in the harp area in Lebinthini specimens. Left panel: male, middle panel: female, right panel: detail of males structures. **A-C**, *Falcerminthus sandakan*; **D-F**, *Agnothecous obscurus*; **G-I**, *Pixibinthus sonicus*; **J-L**, *Ligypterus linarensis*.

**Fig 3:**
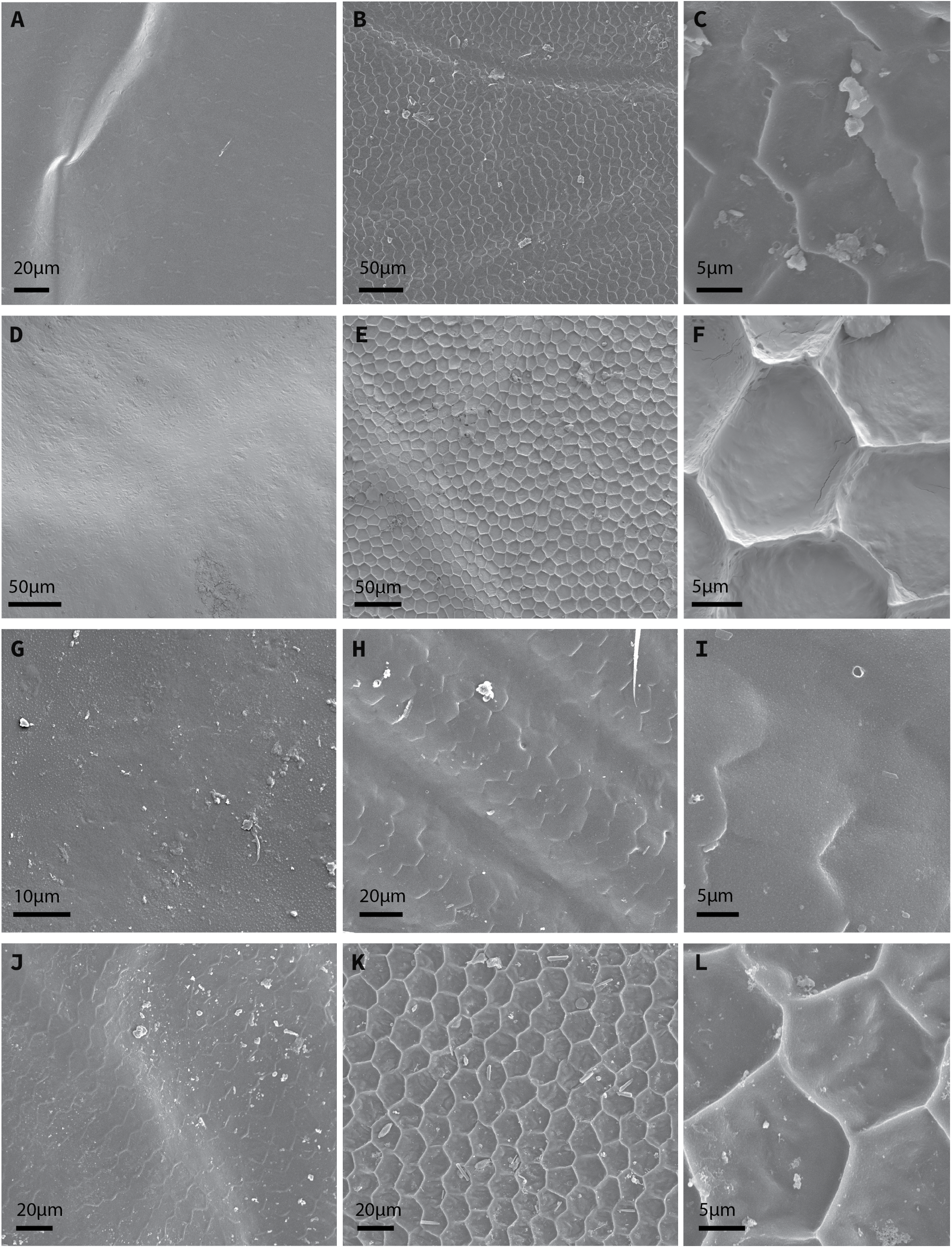
SEM pictures of the dorsal forewings membranes ultrastructure in the harp area in non-Lebinthini specimens. Left panel: male; middle panel: female; right panel: detail of females structures. **A-C**, *Xenogryllus eneopteroides*; **D-F**, *Nisitrus malaya*; **G-I**, *Eneoptera guyanensis*; **J-l**, *Eurepa marginipennis*.

In all the female specimens, the dorsal surface of the forewings membrane is homogeneously covered with hexagonal honeycomb-like or scale-like hexagonal structures, with the structural units (a ‘hexagonal cell’ or a scale) having an average diameter of about

12.5 µm (See Table 2 for species average for each trait). For species of the genera *Centuriarus, Ligypterus* (**Fig. 2J-L**), *Ponca, Eneoptera* (**Fig. 3G-I**) and *Xenogryllus* (**Fig. 3A-C**), the hexagonal cells resemble flat fused cells, without proper walls (all wall thicknesses coded as null) (Table 2).

**Table 1.** Taxonomic sampling. Abbv: M=male, F=female, H=High-frequency, L=low-frequency.

| Tribe | Subtribe | Species | Adults | Wing buds | Call frequency |
| --- | --- | --- | --- | --- | --- |
| Eneopterini |  | *Eneoptera guyanensis* | M+F |  | H |
| Eneopterini |  | *Eneoptera surinamensis* | M+F |  | L |
| Eurepini |  | *Eurepa marginipennis* | M+F |  | L |
| Nisitrini |  | *Nisitrus danum* | M+F | X | L |
| Nisitrini |  | *Nisitrus malaya* | M+F | X | L |
| Nisitrini |  | *Nisitrus vittatus* | M+F |  | L |
| Nisitrini |  | *Nisitrus insignis* | M |  | L |
| Xenogryllini |  | *Pseudolebinthus gorochovi* | M+F |  | H |
| Xenogryllini |  | *Xenogryllus marmoratus* | M+F |  | L |
| Xenogryllini |  | *Xenogryllus eneopteroides* | M+F |  | L |
| Lebinthini | Cardiodactylina | *Cardiodactylus novaeguineae* | M+F |  | H |
| Lebinthini | Cardiodactylina | *Cardiodactylus enkraussi* | M+F |  | H |
| Lebinthini | Cardiodactylina | *Cardiodactylus tankara* | M |  | H |
| Lebinthini | Lebinthina | *Agnothecous obscurus* | M+F |  | H |
| Lebinthini | Lebinthina | *Agnothecous tapinopus* | M+F |  | H |
| Lebinthini | Lebinthina | *Agnothecous azurensis* | M+F |  | H |
| Lebinthini | Lebinthina | *Agnothecous yahoue* | M+F |  | H |
| Lebinthini | Lebinthina | *Centuriarus centurio* | M+F |  | H |
| Lebinthini | Lebinthina | *Falcerminthus sanchezi* | M+F |  | H |
| Lebinthini | Lebinthina | *Falcerminthus sandakan* | M+F | X | H |
| Lebinthini | Lebinthina | *Falcerminthus truncatipennis* | M+F |  | H |
| Lebinthini | Lebinthina | *Falcerminthus villemantae* | M+F |  | H |
| Lebinthini | Lebinthina | *Falcerminthus estrellae* | M+F |  | H |
| Lebinthini | Lebinthina | *Gnominthus baitabagus* | M+F | X | H |
| Lebinthini | Lebinthina | *Lebinthus bitaeniatus* | M+F |  | H |
| Lebinthini | Lebinthina | *Lebinthus luae* | M+F | X | H |
| Lebinthini | Lebinthina | *Macrobinthus jharnae* | M+F | X | H |
| Lebinthini | Lebinthina | *Microbinthus pintaui* | M+F |  | H |
| Lebinthini | Lebinthina | *Microbinthus santoensis* | M+F |  | H |
| Lebinthini | Lebinthina | *Pixibinthus sonicus* | M+F |  | H |
| Lebinthini | Ligypterina | *Ligypterus fuscus* | M+F |  | H |
| Lebinthini | Ligypterina | *Ligypterus linharensis* | M+F |  | H |
| Lebinthini | Ligypterina | *Ponca herbardi* | M+F |  | H |

**Table 2.** Mean values of the traits for each sampled species, with standard error. For density measurements, as all structures in one pictures have been counted, no standard error is available.

| Tribe | Species | Microtrichia density (mic./mm<sup>2</sup>) | | Microtrichia length ($\mu\text{m}$) | | Hexagonal cell diameter ($\mu\text{m}$) | | Hexagonal cell density (mic./mm<sup>2</sup>) | | Hexagonal cell walls thickness ($\mu\text{m}$) | |
| --- | --- | --- | --- | --- | --- | --- | --- | --- | --- | --- | --- |
|  |  | Male | Female | Male | Female | Male | Female | Male | Female | Male | Female |
| Ene. | *E.guyanensis* | 5879 | 5747 | 10.86±3.43 | 12.90±4.78 | 0 | 13.98±2.1 | 0 | 3484 | abs | 0 |
| Ene. | *E.surinamensis* | 5637 | 4474 | 3.34±1.10 | 1.76±0.55 | 0 | 13.87±1.81 | 0 | 6113 | abs | 0 |
| Eur. | *E.marginipennis* | 7098 | 0 | 4.44±1.89 | 0 | 12.76±1.84 | 16±1.57 | 6644 | 4465 | 0 | 0.760±0.146 |
| Nis. | *N.danum* | 4516 | 4452 | 1.07±0.27 | 1.53±0.32 | 0 | 15.03±1.16 | 0 | 4907 | abs | 0.581±0.08 |
| Nis. | *N.malaya* | 6973 | 4516 | 2.20±0.41 | 0.88±0.204 | 0 | 15.55±1.6 | 0 | 4699 | abs | 0.484±0.123 |
| Nis. | *N.vittatus* | 4985 | 3180 | 1.18±0.14 | 2.01±0.56 | 0 | 15.36±1.74 | 0 | 4789 | abs | 0.692±0.142 |
| Nis. | *N.insignis* | 3977 | ? | 2.34±0.56 | ? | 0 | ? | 0 | ? | abs | ? |
| Xen. | *P.gorochovi* | 6167 | 0 | 0.92±0.22 | 0 | 0 | 8.94±1.58 | 0 | 10918 | abs | 0.551±0.222 |
| Xen. | *X.marmoratus* | 5575 | 6964 | 1.61±0.36 | 6.03±1.98 | 0 | 15.50±2.08 | 0 | 4955 | abs | 0 |
| Xen. | *X.eneopteroides* | 5077 | 4907 | 1.34±0.33 | 3.99±1.68 | 0 | 14.27±1.74 | 0 | 5648 | abs | 0 |
| Leb. | *A.obscurus* | 2356 | 0 | 0.72±0.19 | 0 | 11.59±1.59 | 11.04±1.72 | 8640 | 9770 | 0.503±0.11 | 0.615±0.086 |
| Leb. | *A.tapinopus* | 7264 | 0 | 1.47±0.38 | 0 | 12.48±1.33 | 11.51±1.6 | 7836 | 8828 | 0.543±0.12 | 0.440±0.134 |
| Leb. | *A.azurensis* | 9910 | 0 | 1.32±0.27 | 0 | 10.21±1.11 | 9.51±0.98 | 10267 | 12739 | 0.615±0.12 | 0.317±0.129 |
| Leb. | *A.yahoue* | 8850 | 1003 | 1.35±0.29 | 1.02±0.41 | 10.77±1.25 | 11.73±1.48 | 9757 | 8621 | 0.702±0.13 | 0.485±0.154 |
| Leb. | *C.novaeguineae* | 0 | 0 | 0 | 0 | 13.87±1.65 | 15.04±1.98 | 5614 | 4853 | 0.708±0.12 | 0.837±0.137 |
| Leb. | *C.enkraussi* | 2160 | 0 | 1.14±0.26 | 0 | 14.07±1.66 | 14.17±1.95 | 5735 | 6122 | 0.488±0.07 | 0.721±0.182 |
| Leb. | *C.tankara* | 3414 | ? | 1.32±0.49 | ? | 14.39±1.72 | ? | 6132 | ? | 0.544±0.14 | ? |
| Leb. | *C.centuriarius* | 1308 | 0 | 3.36±1.48 | 0 | 14.49±1.46 | 12.98±1.51 | 5609 | 6791 | 0 | 0 |
| Leb. | *F.sanchezi* | 0 | 0 | 0 | 0 | 9.68±1.09 | 8.92±0.91 | 6863 | 14829 | 1.458±0.56 | 0.511±0.209 |
| Leb. | *F.sandakan* | 0 | 0 | 0 | 0 | 11.72±1.28 | 11.55±1.38 | 7605 | 9284 | 2.602±0.48 | 0.497±0.144 |
| Leb. | *F.truncatipennis* | 0 | 0 | 0 | 0 | 12.00±1.31 | 11.54±1.5 | 8521 | 8667 | 0.914±0.20 | 0.623±0.156 |
| Leb. | *F.villemantae* | 0 | 0 | 0 | 0 | 13.31±1.35 | 10.64±1.16 | 6825 | 10443 | 0.820±0.28 | 0.693±0.126 |
| Leb. | *F.estrellae* | 0 | 0 | 0 | 0 | 10.08±1.22 | 10.20±1.07 | 10279 | 11944 | 0.687±0.16 | 0.699±0.215 |
| Leb. | *G.baitabagus* | 623 | 0 | 0.87±0.22 | 0 | 10.52±1.24 | 14.49±1.29 | 9485 | 5803 | 0.599±0.13 | 0.888±0.174 |
| Leb. | *L.bitaeniatus* | 0 | 0 | 0 | 0 | 12.88±1.64 | 12.26±1.55 | 6811 | 7351 | 1.555±0.38 | 0.619±0.151 |
| Leb. | *L.luae* | 0 | 0 | 0 | 0 | 12.83±1.29 | 13.47±1.57 | 7235 | 6244 | 0.834±0.204 | 0.734±0.186 |
| Leb. | *L.fuscus* | 6636 | 0 | 1.03±0.23 | 0 | 11.82±1.48 | 13.27±1.73 | 7219 | 9081 | 0 | 0 |
| Leb. | *L.linarensis* | 7867 | 15607 | 1.47±0.29 | 0.73±0.18 | 13.57±1.57 | 12.75±1.93 | 7457 | 8005 | 0 | 0 |
| Leb. | *M.jharnae* | 1094 | 0 | 0.75±0.17 | 0 | 12.83±1.30 | 11.25±2.05 | 7967 | 6879 | 0.580±0.37 | 0.630±0.109 |
| Leb. | *M.pintaudi* | 14477 | 0 | 1.13±0.30 | 0 | 11.17±1.41 | 10.74±1.1 | 10568 | 9063 | 0.463±0.18 | 0.564±0.12 |
| Leb. | *M.santoensis* | 11476 | 1919 | 1.45±0.36 | 1.75±0.63 | 10.19±1.16 | 11.47±1.41 | 8314 | 9840 | 0.304±0.12 | 0.552±0.113 |
| Leb. | *P.sonicus* | 14515 | 0 | 1.08±0.24 | 0 | 11.17±1.11 | 13.13±1.41 | 8818 | 6424 | 0.525±0.16 | 0.532±0.202 |
| Leb. | *P.hebardi* | 7743 | 0 | 1.06±0.67 | 0 | 10.85±1.54 | 12.49±1.34 | 8249 | 8090 | 0 | 0 |

Male specimens of the genera *Eneoptera, Nisitrus, Xenogryllus* and *Pseudolebinthus* present a completely smooth wing membrane in the harp area, characterized by the absence of hexagonal cell structures (**Fig. 3**). Other areas of the forewings membrane, such as the lateral field, can be covered with hexagonal structures in these genera. For the other species, the hexagonal cells are present in the harp region with an average diameter of ca. 11.9 μm for the males (Table 2). For species of the genera *Centuriarus, Eurepa, Ligypterus* and *Ponca*, as for female specimens, the hexagonal cells are more like flat fused cells, without proper walls (wall thickness considered as null). For the remaining species, i.e. most of the Lebinthini species, the average cell walls thickness in males is about 0.83 µm, with maximum values reaching up to 2.6 µm in *Falcerminthus sandakan*, or about 1.5 µm in *F. sanchezi* and *L. bitaeniatus*.

On the ventral surface, the presence of more or less filamentous microtrichia is noted (**Fig. 4**,**5**). Except for the species of the genera *Falcerminthus* and *Lebinthus*, and *Cardiodactylus novaeguineae*, all the male samples show these microtrichia in the harp region, with an average density of about 6e10^3^ microtrichia per mm^2^ of membrane. The density of microtrichia in the harp region in males ranges from absence of the structures to 14,515 microtrichia per mm^2^ in *Pixibinthus sonicus*. In most of the male Lebinthini samples, the microtrichia are very short, resembling simple knobs (**Fig. 4**), while they can measure up to 11 µm in male *E. guyanensis*. (**Fig. 5G**).

**Fig 4:**
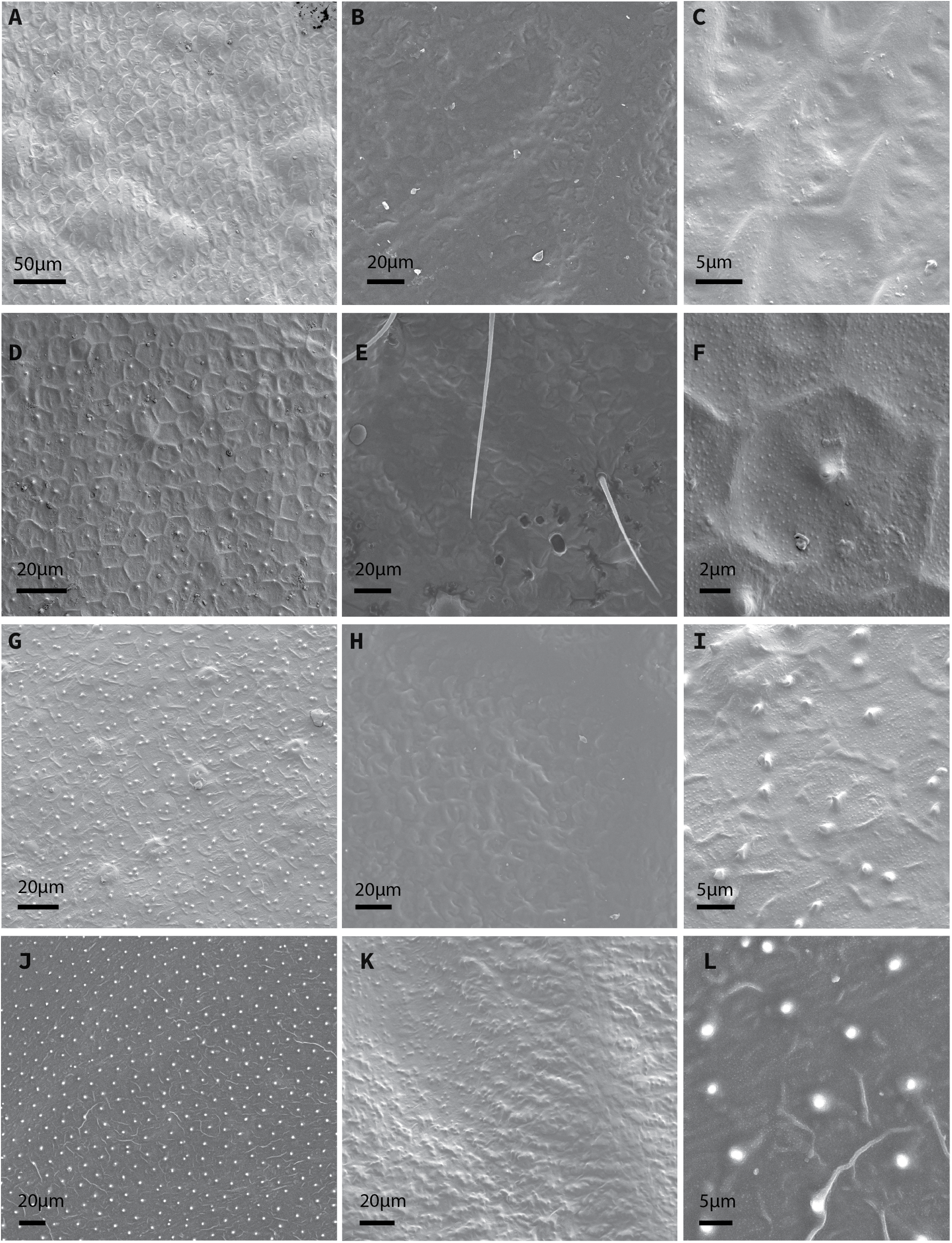
SEM pictures of the ventral face of the forewing membranes ultrastructure in the harp area in Lebinthini specimens. Left panel: male, middle panel: female, right panel: detail of males structures. **A-C**, *Falcerminthus sandakan*; **D-F**, *Agnothecous obscurus*; **G-I**, *Pixibinthus sonicus*; **J-L**, *Ligypterus linarensis*.

**Fig 5:**
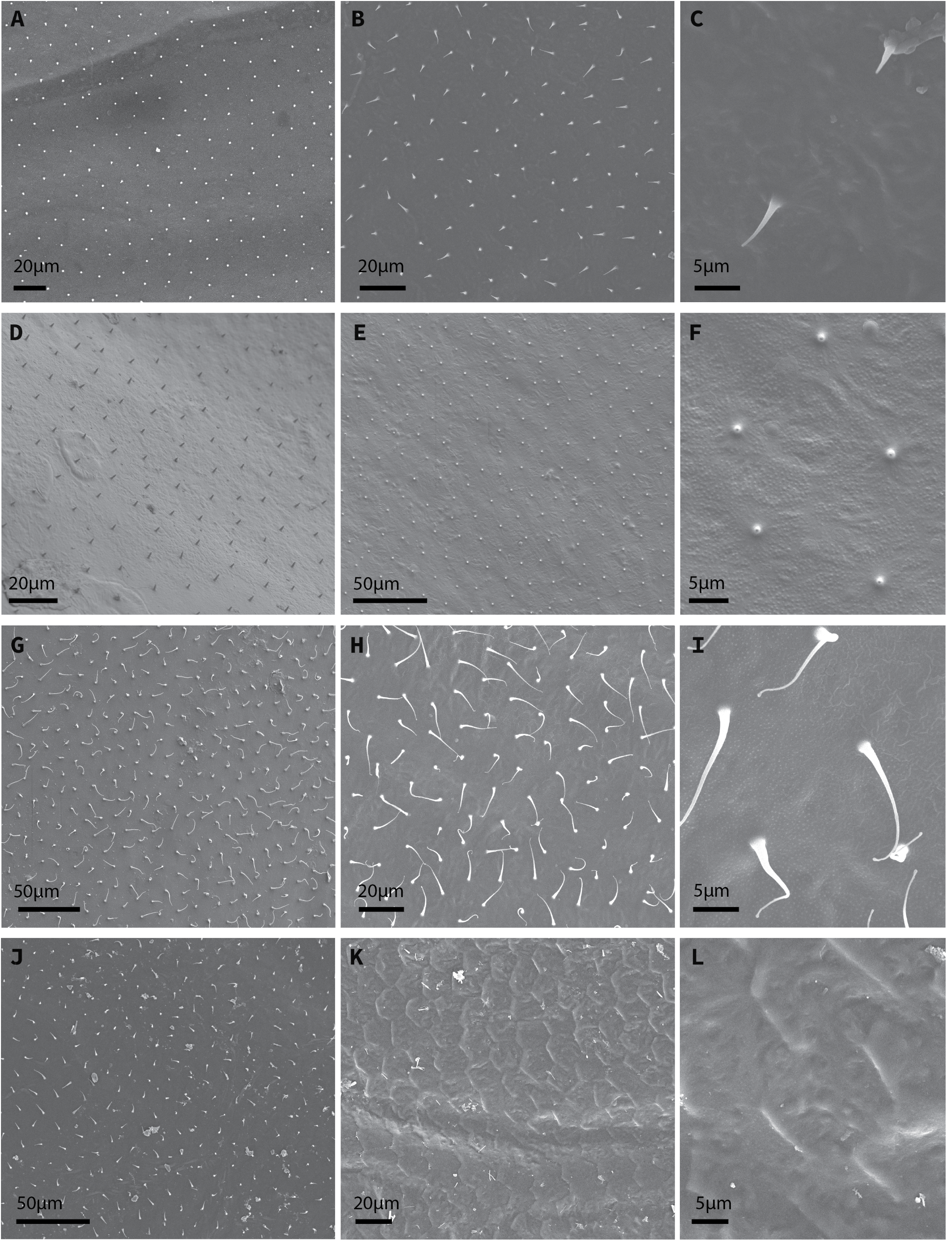
SEM pictures of the ventral forewing membranes ultrastructure in the harp area in non-Lebinthini specimens. Left panel: male; middle panel: female; right panel: detail of females structures. **A-C**, *Xenogryllus eneopteroides*; **D-F**, *Nisitrus malaya*; **G-I**, *Eneoptera guyanensis*; **J-l**, *Eurepa marginipennis*.

The female samples all show microtrichia in small areas in the basal part of the wing, but only 10 out of the 31 female samples show microtrichia on the main surface and the sampled region, among which only 3 Lebinthini species (Table 2). As for the males, the longest microtrichia are observed in *E. guyanensis (*ca. 13 µm) (Table 2).

We also observe other micro-structures on the ventral face (**Fig 4, 5K**) of male and female forewing, resembling attenuated versions of the dorsal hexagonal structures. Because of the diversity in terms of shape within a given sample, no measurements were made about these structures.

Wilcoxon paired samples mean comparison tests (**Fig. 6**) showed significant mean differences between males and females for the whole subfamily for microtrichia density (p.value = 0.00062**), and length (p.value = 0.04635*), and for hexagonal cells density (p.value = 0.02633*) and diameter (p.value = 0.00509**). For the hexagonal cells thickness, the H0 hypothesis of mean equality could not be rejected (p.value = 0.659).

**Fig 6:**
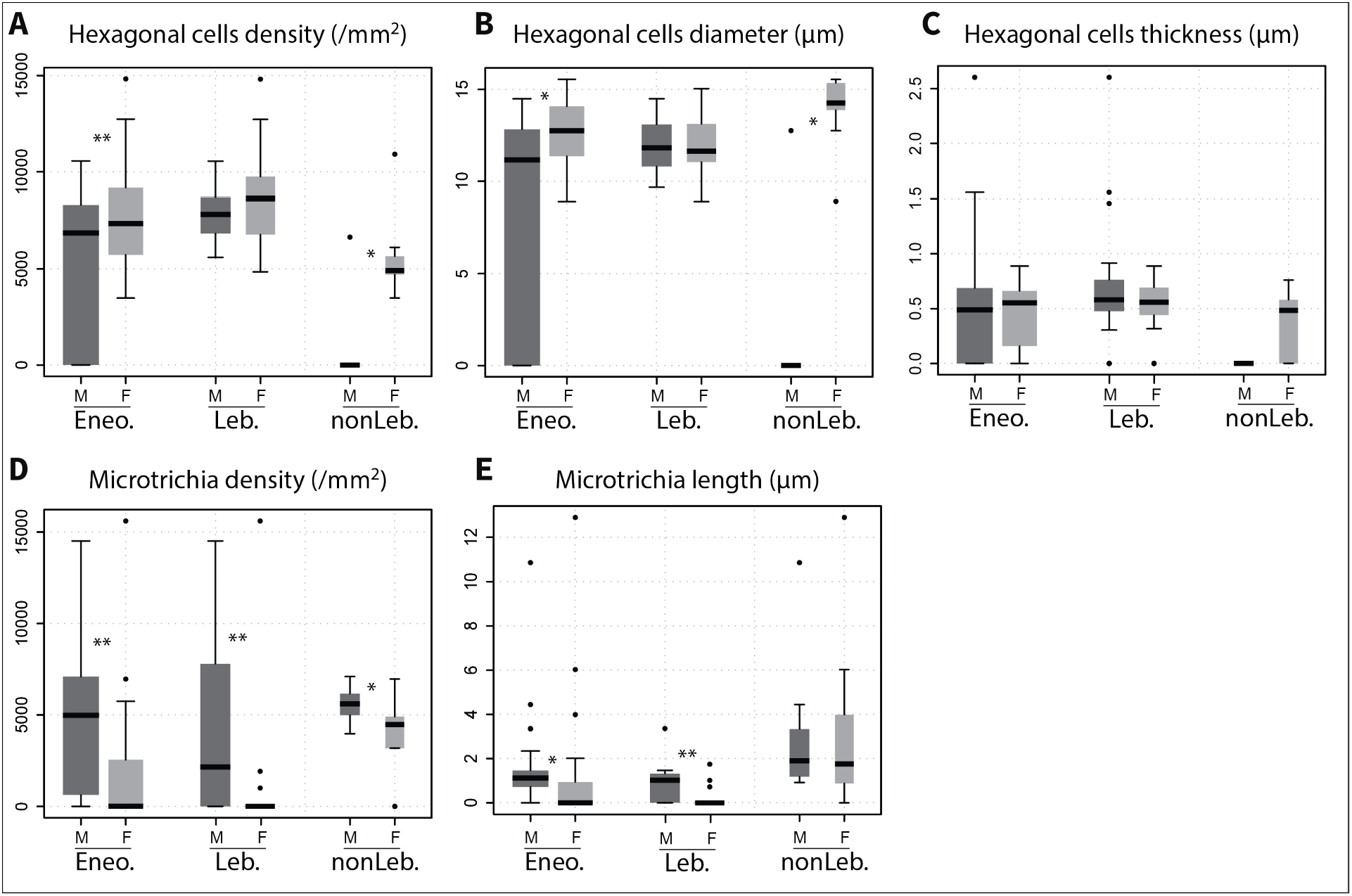
Difference of trait values between males (M) and females (F) in the whole subfamily (Eneo.), in Lebinthini tribe (Leb.) and in non-Lebinthini species (nonLeb.). Stars indicate significant differences between males and females from Wilcoxon paired samples mean comparison tests.

Within the Lebinthini, no more significant difference between males and females is found for hexagonal cells related traits (p.value “hexagon density” = 0.209928; p.value hexagon diameter = 0.949270, hexagonal cells thickness = 0.338008). However, there is still a significant difference between males and females for the microtrichia density (p.value = 0.005742**) and length (p-value = 0.001367**) in the Lebinthini.

For the non-Lebinthini species, there is no more significant difference between males and females for microtrichia length (p-value = 0.910156), but the microtrichia density is still significantly different (p-value = 0.039063*). As for the whole subfamily, hexagonal cells are different between males and females in terms of diameter (p.value = 0.014266*) and density (p-values = 0.007813**). The difference in hexagonal cells thickness is no longer significant for this dataset (p-value = 0.059058).

The mean comparison tests between Lebinthini and non Lebinthini for the 3 traits microtrichia length, hexagonal cells diameter, hexagonal cells thickness show significant differences for the 3 traits with great support (see **Table 3**, all p-values < 10e-8), when comparing values of species among males and females separately.

**Table 3.** Mean comparison tests results between Lebinthini and non-Lebinthini species. Stars indicate mean significant differences.

|  | Statistic | p-value |
| --- | --- | --- |
| Males microtrichia length | 41072.5 | **5.193E-62\*\*** |
| Females microtrichia length | 31897 | **8.002E-107\*\*** |
| Males hexagonal cells diameter | 194095 | **1.389E-108\*\*** |
| Females hexagonal cells diameter | 48324 | **5.548E-28\*\*** |
| Males hexagonal cells thickness | 187950 | **1.198E-100\*\*** |
| Females hexagonal cells thickness | 111554.5 | **1.095E-09\*\*** |

For the correlation tests of the ultrastructural traits with body size and harp size, only the non-null values have been considered. No correlation between our traits and the body size have been found for both sexes (**Table 4**). In males the hexagonal cells diameter is however positively correlated to harp size with a correlation coefficient of 0.68 (p-value = 0.000286**), but the hexagonal cell density is negatively correlated to the harp size, with a coefficient of −0.69 (p-value = 0.000168**). We obtained similar results for females with a correlation coefficient of 0.6 (p-value = 0.000316**) for hexagons diameter and −0.57 (p-value = 0.000903**) for their density.

**Table 4.** Correlation tests results of the ultrastructural traits with size in Eneopterinae. **A:** male samples, **B:** female samples. Stars indicate significant correlation.

| **A/ males** | **Microtrichia density** | **Microtrichia length** | **Hexagonal cells diameter** | **Hexagonal cell density** | **Hexagonal cells thickness** |
| --- | --- | --- | --- | --- | --- |
| Body size | r = -0.331542<br>p-val = 0.105445 | r = 0.091010<br>p-val = 0.665268 | r = 0.276220<br>p-val = 0.191365 | r = -0.096034<br>p-val = 0.655318 | r = -0.197824<br>p-val = 0.416893 |
| Harp size | r = -0.296458<br>p-val = 0.150155 | r = 0.315209<br>p-val = 0.124836 | **r = 0.676257**<br>**p-val = 0.000286\*\*** | **r = -0.694249**<br>**p-val = 0.000168\*\*** | r = -0.046272<br>p-val = 0.850796 |

| **B/ females** | **Microtrichia density** | **Microtrichia length** | **Hexagonal cells diameter** | **Hexagonal cell density** | **Hexagonal cells thickness** |
| --- | --- | --- | --- | --- | --- |
| Body size | r = -0.241634<br>p-val = 0.501221 | r = 0.362653<br>p-val = 0.303054 | r = -0.108135<br>p-val = 0.562569 | r = 0.006913<br>p-val = 0.970556 | r = -0.136273<br>p-val = 0.535257 |
| Harp size | r = 0.115568<br>p-val = 0.750543 | r = 0.412943<br>p-val = 0.235602 | **r = 0.604515**<br>**p-val = 0.000316\*\*** | **r = -0.566047**<br>**p-val = 0.000903\*\*** | r = 0.377189<br>p-val = 0.076020 |

### Wing-buds ultrastructure

For all of the 5 sampled species the membrane ultrastructure of the wing buds looked similar for both sexes and developmental stage (Fig.8). The wing buds are covered, on both ventral and dorsal faces, with short microtrichia, with an average density of 1.06E+05 per mm2. No hexagonal structures are visible on any face. In some species on the ventral face, microtrichia seem to be non-randomly arranged (Fig 8M-P), especially for *G. baitabagus*, whose microtrichia are shorter and could sometimes be mistaken with emerging scales.

## Discussion

Our study reveals that there is a great diversity in micro-structures configurations on the dorsal and ventral face of the forewing membrane among the crickets of the Eneopterinae subfamily. Two types of structures are observed; hexagonal cells on the dorsal face of the forewing, and microtrichia on the ventral face, the same structures that have been observed in other cricket families or subfamilies such as Phalangopsidae, Oecanthidae or Gryllinae (Barranco & Molina-Pardo, 2021; Desutter-Grandcolas, 1995). We remark an original diversity in terms of configuration of these structures. In some species these structures can be absent or vary in density or size, among the subfamily. There is also variety of ultrastructure along the wing, with the structures being absent in some areas and present in others, as previously shown by Desutter-Grancolas (1995). For some species, males and females samples exhibit very dissimilar membranes, while other species show little dimorphism. Regarding the different functions of the wing, especially sound production in males, this diversity has to be addressed in a comparative and phylogenetic context.

Eneopterinae females always show hexagonal structures, more or less strong, when the majority of the non-Lebinthini males present a flat smooth membrane. The difference in density and size of these structures is significant between males and females at the scale of the whole subfamily, but this is because of the strong sexual dimorphism for this trait in non-Lebinthini species. In the Lebinthini tribe, males and females of the same species are not significantly different for these traits. We also have a strong difference in ultrastructure for all ultrastructural traits between Lebinthini and non-lebinthini males. The non-Lebinthini species exhibit a flat harp membrane, when the Lebinthini species, who are all producing high-frequency signals associated with harmonic hopping events, still present hexagonal cells similar to those visible in females. Compared to what is observed in other cricket groups, low-frequency singers (non-Enopterinae and non-Lebinthini Eneopterinae) exhibit more flattened structures in the harp region. These results associated with the sexual dimorphism of the trait suggest a link between hexagonal ultrastructures and sound production. We propose the hypothesis that a reduction or loss of the hexagonal structures in males results in making the wing membrane thinner, which allows the wing to vibrate at relatively low frequency.

The present results lead us to another hypothesis: Lebinthini males, all producing high-frequency songs, instead of a thinner harp region, exhibit maintained or exagerated hexagonal structures that could thicken and stiffen the membrane, increasing its resonance frequency (Bennet-Clark, 1999b, Gaiddon et al, in prep). In the species *Lerneca fucipennis* (Phalangopsidae), however, the forewings exhibit hexagons similar to those of the Lebinthini in the harp region, which is in contradion with our hypothesis. But although *L. fuscipennis* is not producing high-frequency calls, it has a particular song described as having frequency modulations (Desutter-Grandcolas, 1998).

Particular ultrastructures are also reported among Lebinthini males, such as thickened hexagons in *Falcerminthus* and *Lebinthus*, or dorsal microtrichia in *Pixibinthus sonicus*. They could potentially be linked to further diversification in frequency value in the tribe, as song can be ultrasonic in some clades or correspond to different harmonic peaks (Tan et al., 2021).

The precedent hypothesis should be tested in a phylogenetic framework using phylogenetic comparative analysis. Additional information of the non-Eneopterinae female structures would be necessary to any further conclusions.

Microtrichia, the second ultrastructure type present on the ventral face, are not evenly distributed between males and females in all tribes, but the difference in length is no longer significant for non-Lebinthini species. Significant difference between lebinthini and non-Lebinthini is found for both density and length. However these structures could be completely independent of the vibratory properties of the harp membrane. In many insects microtrichia are sensory structures involved in the perception of the position of the wings against the body or against each other (Gorb, 1999; Sun et al., 2018). The very small size of these structures in Lebinthini (reduced to knobs) may cast doubt on their sensory role, in contrast to what is observed in some species where microtriches are very long and likely to come into contact with bristle-type sensory sensors (the microtriches themselves being devoid of nerve endings) (Czaja, 2013; Desutter-Grandcolas, 1995; Gorb, 1999; Qian et al., 2016). In females, the absence or great reduction of microtrichia could be related to the absence of use of the forewing (no singing nor flight). This hypothesis is supported in Lebinthini by the fact that female forewings tend to be reduced and to become so small they no longer overlap.

The five measured traits are not correlated to body size, but hexagon’s size and density is related to harp size in both males and females of the subfamily. The hexagonal cell diameter is positively correlated to harp size, when the density is negatively correlated to the harp size. The size-effect is important in song production (Montealegre-Z, 2009) and thus has to be considered for further analyses testing the potential correlation between these traits and acoustic features.

In juveniles, the hexagonal structures on the dorsal surface of adults are not yet visible on the wing buds, whatever the stage, suggesting these structures to be put in place during the last imaginal molt. In Orthoptera (including crickets), between the last juvenile stage and the adult, the forewings undergo an inversion, so the ventral face of the wing bud corresponds to what will be the dorsal face of the adult wing. On this ventral side of the buds, in the three Lebinthini species sampled (Fig. 7A-C, G-I, M-P), the microtrichia appear to be arranged like scales in both sexes. This could suggest that the formation of hexagonal ultrastructures walls during development may originate from fused microtrichia. However, *N. malaya* females’ wing buds microtrichia do not show such an arrangement, whereas adult females do have very pronounced hexagonal structures. More study on such larval structures in non-Eneopterinae crickets could bring light on the development of the adult microstructures.

**Fig 7:**
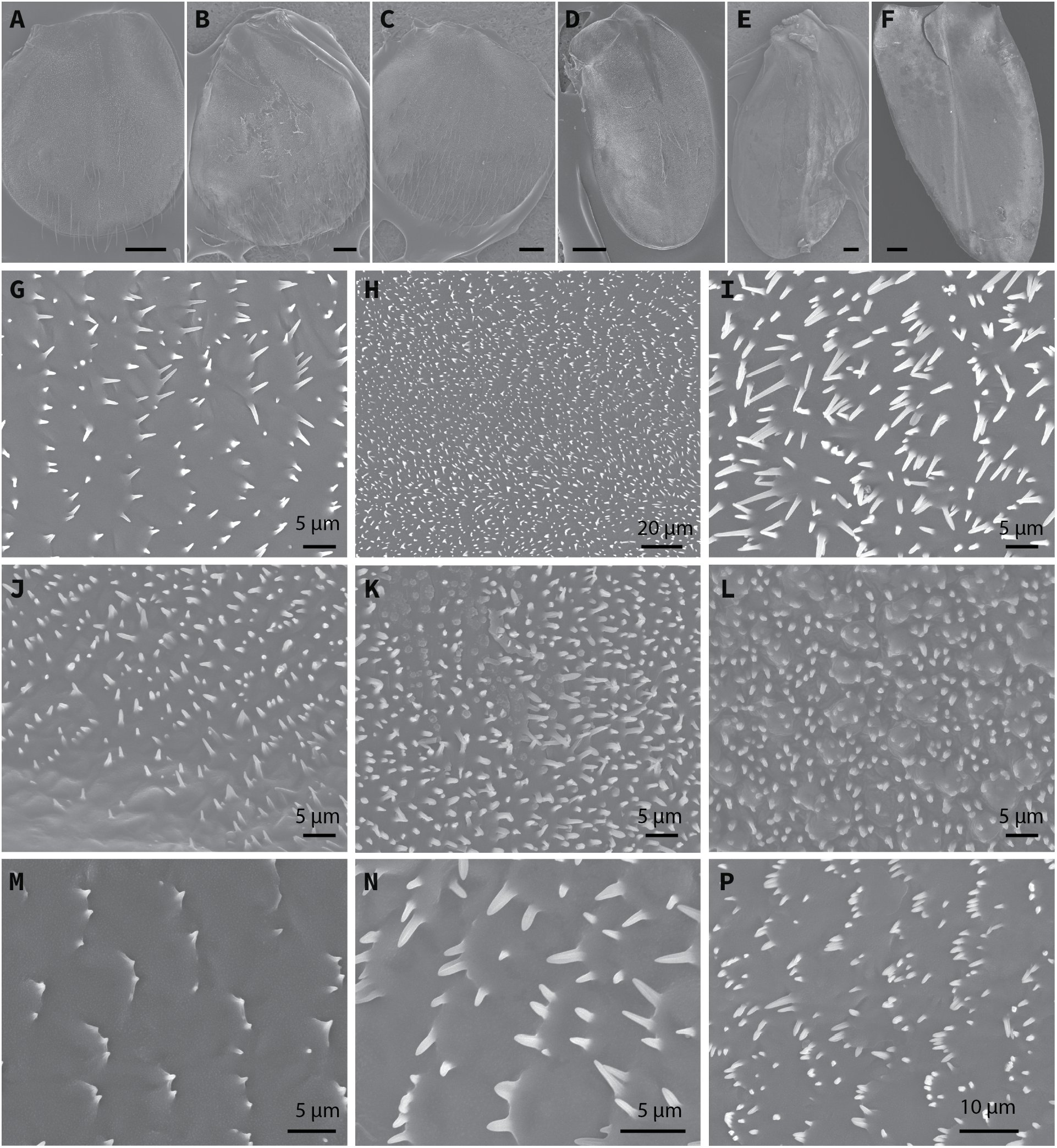
SEM pictures of wing bud ultrastructures in Enopterinae specimens. **Top panel:** Dorsal overviews of wing buds. Scale bars = 200µm **A-C** *Lebinthus luae*. **A:** male first wing bud, **B:** male second wing bud, **C:** female second wing bud. **D-F** *Nisitrus danum*. **D:** male first wing bud, **E:** male second wing bud, **F:** female second wing bud. **G-I** *Lebinthus luae*. **G:** male first wing bud, ventral face, **H:** male second wing bud, ventral face, **I:** male second wing bud, dorsal face, **J-L** *Nisitrus danum*. **J:** male first wing bud, ventral face, **K:** male first wing bud, dorsal face, **L:** male second wing bud, dorsal face. **M-P** Ventral face. **M:** *Gnominthus baitabagus* female second wing bud, **N:** *Macrobinthus jharnae* male first wing bud, **P:** *Macrobinthus jharnae* female second wing bud.

## Conclusions

The present study highlights significant diversity in ultrastructural configuration for both dorsal hexagonal cells and ventral microtrichia, with notable variations in their presence, density, and size among species and sexes in the Eneopterinae subfamily. This diversity could be closely linked to sound production in males. The presence of pronounced hexagonal cells in Lebinthini males, in contrast to the flattened harp membrane of non-Lebinthini species, supports the hypothesis that these structures enhance resonance frequency in the harp region, resulting in high-frequency songs with harmonic hopping. Further comparative and phylogenetic analyses are essential to explore these traits’ roles in biomechanics and evolution comprehensively.

## References

Barranco, P., & Molina-Pardo, J. L. (2021). Cuticular Structures in Micropterous Crickets (Orthoptera, Gryllidae, Petaloptilini, Gryllomorphini). Insects, 12(8), Article 8. 10.3390/insects12080708

Bennet-Clark, H. C. (1999a). Resonators in insect sound production. 11.

Bennet-Clark, H. C. (1999b). Resonators in insect sound production: How insects produce loud pure-tone songs. Journal of Experimental Biology, 202(23), 3347–3357. 10.1242/jeb.202.23.3347

Bradbury, J. W., & Vehrencamp, S. L. (2011). Principles of animal communication, 2nd ed (pp. xiv, 697). Sinauer Associates.

Clark, C. J. (2014). Harmonic Hopping, and Both Punctuated and Gradual Evolution of Acoustic Characters in Selasphorus Hummingbird Tail-Feathers. PLOS ONE, 9(4), e93829. 10.1371/journal.pone.0093829

Czaja, J. (2013). Microtrichial patterns of the mesothoracic wing surface in Scutelleridae (Hemiptera). The Canadian Entomologist, 145(4), 351–368. 10.4039/tce.2013.22

Desutter-Grandcolas, L. (1995). Functional forewing morphology and stridulation in crickets (Orthoptera, Grylloidea). Journal of Zoology, 236(2), 243–252. 10.1111/j.1469-7998.1995.tb04491.x

Desutter-Grandcolas, L. (1998). Broad-frequency modulation in cricket (Orthoptera, Grylloidea) calling songs: Two convergent cases and a functional hypothesis. Canadian Journal of Zoology, 76(12), 2148–2163. 10.1139/z98-152

Eisenbeis, G., & Wichard, W. (2012). Atlas on the Biology of Soil Arthropods. Springer Science & Business Media.

Gerhardt, H. C., & Huber, F. (2002). Acoustic Communication in Insects and Anurans: Common Problems and Diverse Solutions. University of Chicago Press.

Gorb, S. N. (1999). Ultrastructure of the thoracic dorso-medial field (TDM) in the elytra-to-body arresting mechanism in Tenebrionid Beetles (Coleoptera: Tenebrionidae). Journal of Morphology, 240(2), 101–113. 10.1002/(SICI)1097-4687(199905)240:2<101::AID-JMOR2>3.0.CO;2-7

Hofstede, H. M. ter, Schöneich, S., Robillard, T., & Hedwig, B. (2015). Evolution of a Communication System by Sensory Exploitation of Startle Behavior. Current Biology, 25(24), 3245–3252. 10.1016/j.cub.2015.10.064

Kingston, T., & Rossiter, S. J. (2004). Harmonic-hopping in Wallacea’s bats. Nature, 429(6992), Article 6992. 10.1038/nature02487

Li, X., Zhang, X., Luo, W., Wang, Y., & Ren, B. (2016). Micromorphological differentiation of left and right stridulatory apparatus in crickets (Orthoptera: Gryllidae). Zootaxa, 4127(3), 553. 10.11646/zootaxa.4127.3.8

Michels, J., Appel, E., & Gorb, S. N. (2016). Functional diversity of resilin in Arthropoda. Beilstein Journal of Nanotechnology, 7, 1241–1259. 10.3762/bjnano.7.115

Michelsen, A. (1998). The Tuned Cricket. Physiology, 13(1), 32–38. 10.1152/physiologyonline.1998.13.1.32

Montealegre-Z, F. (2009). Scale effects and constraints for sound production in katydids (Orthoptera: Tettigoniidae): correlated evolution between morphology and signal parameters. Journal of Evolutionary Biology, 22(2), 355–366. 10.1111/j.1420-9101.2008.01652.x

Montealegre-Z, F., Jonsson, T., & Robert, D. (2011). Sound radiation and wing mechanics in stridulating field crickets (Orthoptera: Gryllidae). Journal of Experimental Biology, 214(12), 2105–2117. 10.1242/jeb.056283

Qian, J., Chi, D., & Chai, R. (2016). Possible functions of the microtrichia on the cuticle of Ulomoides dermestoides (Chevrolat) (Coleoptera: Tenebrionidae). Journal of Forestry Research, 27(6), 1391–1405. 10.1007/s11676-016-0261-y

R Core Team, R. F. for S. C. (2022). R: The R Project for Statistical Computing. https://www.r-project.org/

Robillard, T., & Desutter-Grandcolas, L. (2004). Phylogeny and the modalities of acoustic diversification in extant Eneopterinae (Insecta, Orthoptera, Grylloidea, Eneopteridae). Cladistics, 20(3), 271–293. 10.1111/j.1096-0031.2004.00025.x

Robillard, T., Grandcolas, P., & Desutter-Grandcolas, L. (2007). A shift toward harmonics for high-frequency calling shown with phylogenetic study of frequency spectra in Eneopterinae crickets (Orthoptera, Grylloidea, Eneopteridae). Canadian Journal of Zoology, 85(12), 1264–1275. 10.1139/Z07-106

Robillard, T., Montealegre-Z, F., Desutter-Grandcolas, L., Grandcolas, P., & Robert, D. (2013). Mechanisms of high-frequency song generation in brachypterous crickets and the role of ghost frequencies. Journal of Experimental Biology, 216(11), 2001–2011. 10.1242/jeb.083964

Robillard, T., ter Hofstede, H. M., Orivel, J., & Vicente, N. M. (2015). Bioacoustics of the Neotropical Eneopterinae (Orthoptera, Grylloidea, Gryllidae). Bioacoustics, 24(2), 123–143. 10.1080/09524622.2014.996915

Sample, C. S., Xu, A. K., Swartz, S. M., & Gibson, L. J. (2015). Nanomechanical properties of wing membrane layers in the house cricket (Acheta domesticus Linnaeus). Journal of Insect Physiology, 74, 10–15. 10.1016/j.jinsphys.2015.01.013

Schindelin, J., Arganda-Carreras, I., Frise, E., Kaynig, V., Longair, M., Pietzsch, T., Preibisch, S., Rueden, C., Saalfeld, S., Schmid, B., Tinevez, J.-Y., White, D. J., Hartenstein, V., Eliceiri, K., Tomancak, P., & Cardona, A. (2012). Fiji: An open-source platform for biological-image analysis. Nature Methods, 9(7), Article 7. 10.1038/nmeth.2019

Sun, J., Liu, C., Bhushan, B., Wu, W., & Tong, J. (2018). Effect of microtrichia on the interlocking mechanism in the Asian ladybeetle, Harmonia axyridis (Coleoptera: Coccinellidae). Beilstein Journal of Nanotechnology, 9(1), 812–823. 10.3762/bjnano.9.75

Tan, M. K., Malem, J., Legendre, F., Dong, J., Baroga-Barbecho, J. B., Yap, S. A., Wahab, R. bin H. A., Japir, R., Chung, A. Y. C., & Robillard, T. (2021). Phylogeny, systematics and evolution of calling songs of the Lebinthini crickets (Orthoptera, Grylloidea, Eneopterinae), with description of two new genera. Systematic Entomology, 46(4), 1060–1087. 10.1111/syen.12510

Vicente, N., Kergoat, G. J., Dong, J., Yotoko, K., Legendre, F., Nattier, R., & Robillard, T. (2017). In and out of the Neotropics: Historical biogeography of Eneopterinae crickets. Journal of Biogeography, 44(10), 2199–2210. 10.1111/jbi.13026

